# Fungal volatiles drive lifestyle-dependent, systemic metabolic reprogramming in poplar

**DOI:** 10.64898/2026.01.19.700287

**Authors:** Peiyuan Zhu, Ina Zimmer, Prasath Balaji Sivaprakasam Padmanaban, Maaria Rosenkranz, Andrea Ghirardo, Jörg-Peter Schnitzler

**Affiliations:** Research Unit Environmental Simulation, Helmholtz Munich, Neuherberg, Germany; Institute of Plant Sciences, Ecology and Conservation Biology, University of Regensburg, Regensburg, Germany

**Keywords:** volatile organic compounds, fungal volatiles, plant-fungal communication, metabolomics, systemic signalling, *Populus*, root-to-shoot communication

## Abstract

Rationale: Although trees encounter diverse fungal communities, it is unclear how they adjust their physiology in response to fungal ecological strategies before physical contact. We tested whether volatile organic compounds (VOCs) emitted by fungi are sufficient to induce systemic, lifestyle-consistent metabolic states in poplar roots and leaves.
Methods: *Populus* × *canescens* roots were exposed to VOCs from a pathogen (*Heterobasidion annosum*), a saprotroph (*Postia placenta*) or an ectomycorrhizal mutualist (*Laccaria bicolor*) for six weeks in a contact-free pot-in-pot system. Untargeted LC–MS metabolomics characterized VOC-induced metabolic reprogramming in roots and leaves.
Key results: Fungal VOC exposure alone reconfigured the metabolomes of roots and leaves, with strong discrimination between treatments despite belowground exposure. Poplar revealed a shared VOC-responsive component, but also fungus-specific programmes: pathogen VOCs produced a suppression-dominated systemic phenotype; saprotroph VOCs promoted lipid-centred metabolic activation; and mutualist VOCs elicited restrained, compatibility-consistent shifts with targeted pathway modulation.
Main conclusion: Volatile-mediated surveillance allows trees to anticipate fungal lifestyle-associated cues and adjust systemic metabolism before physical contact occurs. This links airborne fungal cues to whole-plant physiological configuration and extends plant–fungal recognition beyond contact-dependent mechanisms.

**One-sentence summary:** Volatile-mediated surveillance allows trees to anticipate fungal lifestyle and adjust systemic metabolism before physical contact occurs.

## Introduction

Plants develop and function within dense belowground communities, where their roots are continuously exposed to fungi that span pathogenic, saprotrophic, and mutualistic lifestyles. These interactions influence nutrient acquisition, growth, and stress resistance, and are particularly important in forest ecosystems, where long-lived hosts must process microbial signals over extended periods (Philippot *et al*., 2013; van der Heijden *et al*., 2015; Baldrian, 2017). Although contact-dependent recognition, such as pattern-recognition receptors detecting fungal cell-wall components and triggering immune signalling, is well described (Spoel & Dong, 2012; Doehlemann *et al*., 2017), the manner in which plants assess fungal identity and interaction context prior to physical contact remains unclear.

Volatile organic compounds (VOCs) offer a plausible route for such pre-contact recognition. VOCs are small, diffusible metabolites that move through soil, air and water, enabling information transfer between spatially separated organisms (Peñuelas *et al*., 2014; Weisskopf *et al*., 2021). Fungal volatilomes are chemically diverse and responsive to physiology and the environment. Reviews have highlighted their ecological roles as mediators of interspecific interactions (Morath *et al*., 2012; Yin *et al*., 2025). Fungal VOC blends are shaped by trophic mode and ecological strategy, suggesting that plants may extract lifestyle information from the structure of blends rather than from individual compounds (Müller *et al*., 2013; Guo *et al*., 2021).

The concept of chemical communication displays (CCDs) is an explicitly mixture-based view, capturing multicomponent bouquets that encode biologically meaningful information in their composition, relative ratios, and covariation among components (Junker *et al*., 2018). Guo et al. (2021) applied the CCD theory to fungi, demonstrating that VOC patterns can predict trophic mode and lifestyle across taxa, and quantifying ’phenotypic integration’ (compound covariation) within volatilomes. The CCD theory thus offers a mechanistic framework for how plants could decode ’who is there’ from volatile bouquets, even when individual components vary with growth stage or substrate.

Experimental evidence supports volatile-mediated plant–fungal interactions. Fungal VOCs can alter plant growth, root system architecture, and defence-related physiology without hyphal contact (Bitas *et al*., 2013; Moisan *et al*., 2020; Singh *et al*., 2021), which aligns with the broader role of microbe-induced plant volatiles in plant-microbe ecology (Sharifi & Ryu, 2018). Sesquiterpenes emitted by the ectomycorrhizal fungus *Laccaria bicolor* (Maire) P.D. Orton are sufficient to reprogramme host root development in the absence of physical interaction, demonstrating that VOCs can act as targeted developmental signals (Ditengou *et al*., 2015). However, many studies focus on short-term responses or individual compounds, leaving unresolved the question of whether plants can reliably discriminate among functionally distinct fungal lifestyles based on complex VOC mixtures under standardised conditions.

Using a contact-free pot-in-pot system, we recently reported in Zhu *et al*. (2025, preprint) that Grey poplar (*Populus* × *canescens* (Aiton) Sm.) can discriminate VOC cues from three fungi: a pathogenic fungus (*Heterobasidion annosum* (Fr.) Bref.), a saprotrophic fungus (*Postia placenta* (Fr.) Niemelä, K.H. Larss. & Schigel), and an ectomycorrhizal fungus (*L. bicolor*). Root-zone headspace VOC signatures were dominated by sesquiterpenes and remained stable and lifestyle-specific over time. In contrast, leaf VOC emissions were more diverse and partially converged (Zhu *et al*., 2025, preprint). While these findings establish volatile-mediated fungal lifestyle recognition as a robust phenomenon in a woody host, they do not explain how volatile cues are translated into internal, systemic physiological states.

Plant hormone signalling networks, particularly those involving jasmonates (JA), ethylene (ET) and salicylic acid (SA), provide a mechanistic context for such state transitions (Spoel & Dong, 2012; Wasternack & Hause, 2013). In antagonistic interactions, early JA/ET-linked responses often precede or interact with SA-associated systemic signalling, whereas mutualisms frequently involve active modulation rather than straightforward activation of these pathways. In the *Populus*-*L. bicolor* symbiosis, jasmonate signalling is attenuated by fungal effectors such as MiSSP7, which interfere with JAZ-mediated regulation, suppressing defence and permitting developmental reprogramming (Felten *et al*., 2009; Daguerre *et al*., 2020). This context-dependent adjustment is evolutionarily significant as ectomycorrhizal fungi have evolved repeatedly from saprotrophic ancestors, accompanied by the extensive remodelling of capacities related to decay and the emergence of traits associated with symbiosis (Kohler *et al*., 2015; Wu *et al*., 2022).

Here, we investigate whether the distinct fungal volatile bouquets produced by lifestyle-structured fungi are translated into distinct systemic metabolic programmes in woody hosts. We analysed the metabolomes of the roots and leaves of Grey poplar, which were exposed to VOCs from *H. annosum*, *P. placenta* or *L. bicolor* for six weeks, using the same contact-free exposure experiment and plant cohort as in Zhu et al. (2025, preprint) for volatile profiling. Integrating endpoint metabolomics with time-resolved volatilomics and CCD theory (Junker *et al*., 2018; Guo *et al*., 2021) allows us to test the hypothesis that fungal VOCs act as early lifestyle cues, which are decoded by the roots and integrated into hormone-weighted whole-plant metabolic states prior to physical contact.

## Materials and Methods

### Plant material and growth conditions

Wild-type *Populus* × *canescens* (Aiton) Sm. (INRA clone 717-1B4, Mader *et al.,* 2016) plantlets were used. Plantlets from sterile stock were sectioned and cultured in glass jars with half-strength Murashige & Skoog medium (Duchefa Biochemie, Haarlem, Netherlands) in a growth chamber at 20/16 ℃ (day/night) with a 10/14 h photoperiod (100 µmol m⁻² s⁻¹ PPFD) for two months. Rooted plantlets were transferred to autoclaved substrate (soil: vermiculite: perlite, 3:1:1, v/v/v) and acclimated in a phytochamber at 22/16 ℃ (light/dark), 60/80% relative humidity (light/dark), and a 10/14 h photoperiod (150 µmol m⁻² s⁻¹ PPFD) for three months, with gradual removal of plastic covers over two weeks.

### Fungal material and culture conditions

Three fungal strains representing distinct ecological lifestyles were applied: pathogenic *Heterobasidion annosum* (Fr.) Bref. (stock: Karin Pritsch, Helmholtz Munich), ectomycorrhizal *Laccaria bicolor* (Maire) P.D. Orton (S238N), and saprotrophic *Postia placenta* (Fr.) Niemelä, K.H. Larss. & Schigel (stock: K. Pritsch). Cultures were grown in 35 mm × 10 mm dishes (Corning 430588, Corning Inc., NY, USA) on modified Melin-Norkrans (MMN) medium (1% (w/v) glucose, 0.25% (w/v) NH₄-tartrate, 0.025% (w/v) (NH₄)₂SO₄, 0.05% (w/v) KH₂PO₄, 0.015% (w/v) MgSO₄·7H₂O, 0.005% (w/v) CaCl₂·2H₂O, 0.0025% (w/v) NaCl, 0.01% (w/v) FeCl₃, 0.001% (w/v) thiamine hydrochloride, 1% (w/v) Gelrite; pH 5.2; Guo et al., 2021). Fungal cultures were maintained at room temperature in the dark.

### Pot-in-pot exposure system

A custom pot-in-pot (PiP) system enabled root exposure to fungal VOCs without physical contact (for details see also Zhu et al., 2025, preprint). Upper pots (180 g autoclaved substrate) contained perforations in the lower 2 cm (28 holes, 1 mm diameter) to allow gas exchange and were positioned above lower pots containing fungal cultures. Poplar-present systems were assigned to four lower-pot treatments: fungus-free MMN control, *H. annosum*, *L. bicolor*, and *P. placenta* (VOC exposure: n = 6 plants per treatment). Trays were rotated every three days. Fungal plates were replaced weekly with pre-colonised cultures (*H. annosum*: 8 d; *L. bicolor*: 28 d; *P. placenta*: 4 d pre-cultivation), and control systems received fresh fungus-free MMN plates on the same schedule. VOC exposure lasted six weeks. This pot-in-pot exposure experiment is the same experimental run used for volatile profiling in Zhu et al. (2025, preprint). Specifically, the VOC exposure design (including fungal pre-cultivation times and weekly plate replacement), growth conditions, and tray rotation scheme were identical, and the root and leaf tissues used for metabolomics in the present study were collected from those same plants at the end of the six-week exposure period.

### Sample collection and metabolite extraction

Plant harvest for the metabolomic analysis coincided with the final volatile sampling (described in detail in Zhu *et al.,* 2025, preprint), enabling direct comparison of VOC signatures and metabolome outcomes within the same exposure system. After six weeks of VOC exposure, root and leaf tissues were harvested (n = 4 per treatment; n = 3 for leaf controls). During the VOC exposure, we weekly checked that there was no physical contact between roots and fungi. Roots were excavated, washed with deionized water, thoroughly dried, and flash-frozen in liquid nitrogen. The top leaves (2-3) were pooled per plant and flash frozen. All samples were stored at -80 ℃, then freeze-dried and ground to a fine powder for metabolite extraction, following Bertić *et al*. (2021) and Sivaprakasam Padmanaban *et al*. (2025). Fifty mg of tissue powder was extracted with 1 mL ice-cold methanol/2-propanol/water (1:1:1, v/v/v), vortexed for 30 s, sonicated for 10 min at 4 ℃, and centrifuged at 10,000 × g for 10 min at 4 ℃. Supernatants were vacuum-dried and reconstituted in 200 μL of 50% acetonitrile. Quality control (QC) samples were prepared by pooling equal volumes of all study extracts. Pooled QC injections were analysed every five study samples throughout the sequence to monitor analytical stability. Solvent blanks were injected at the beginning and end of the batch to assess background and carryover.

### Liquid chromatography-mass spectrometry (LC-MS) analysis

Untargeted metabolomics analysis was performed using an ultra-high performance liquid chromatography (UHPLC/UPLC) system coupled to an ultra-high resolution (UHR) QqToF mass spectrometer. Metabolite profiling was conducted on an Ultimate 3000RS UPLC (Thermo Fisher Scientific, Bremen, Germany) coupled to a Bruker Impact II QqToF mass spectrometer equipped with an electrospray ionisation (ESI) source. Chromatographic separation was achieved using reversed-phase LC (RPLC) on an Acquity BEH C18 column (150 mm × 2.1 mm, 1.7 μm; Waters) maintained at 40 ℃, with mobile phases of (A) water + 0.1% (v/v) formic acid and (B) acetonitrile + 0.1% (v/v) formic acid. The injection volume was 5 μL at a flow rate of 0.35 mL min⁻¹ (ESI+) and 0.4 mL min⁻¹ (ESI−). Mass spectrometric detection was conducted in separate runs in both ESI+ and ESI− modes. Key parameters were: capillary voltage 4.5 kV (ESI+) / 3.5 kV (ESI−), end plate offset −500 V, nebulizer pressure 2.0 bar, drying gas 8.0 L min⁻¹ at 200 ℃. Spectra were acquired over m/z 20–1000 at 8 Hz. Data-dependent MS/MS was performed on the top 3 precursors per cycle using CID (collision-induced dissociation).

Raw LC-MS data were processed in MetaboScape 4.0 (Bruker Daltonics, Bremen, Germany) for peak detection, chromatographic alignment, and isotope/adduct grouping. Mass features detected in ESI+ and ESI− modes were processed separately and subsequently merged for downstream analysis. Metabolite annotation was performed based on accurate mass and MS/MS spectral matching following the workflow described by Bertić et al. (2021). Compound identification relied on spectral library searches against databases including KEGG, HMDB, LIPID MAPS, and PlantCyc. Features were reported as MSI level 2 (putatively annotated compounds) when MS/MS matches achieved a spectral score >700 with a mass error <5 ppm. For features without MS/MS matches, elemental formulas were predicted using SmartFormula, and tentative annotations were assigned at MS1 level using 10 mDa tolerance for the precursor mass with in-house R code. All detected features were additionally classified using multidimensional stoichiometric compound classification (MSCC) (Rivas-Ubach *et al*., 2018) based on an assessment of the C/H/O/N/P stoichiometric ratios, enabling functional grouping of unannotated metabolites into biochemical classes (e.g., lipids, secondary metabolites, peptides/amino acids).

### Statistical analysis

Prior to statistical analysis, peak areas were normalized using median normalization to correct for systematic variation among samples. Multivariate analyses on mass feature data were performed in SIMCA 13 (v13.0.3.0, Umetrics, Umeå, Sweden) and univariate statistics and visualisation in R v4.3.1 (R Core Team, 2023). Before multivariate analyses, mass feature data were log_10_-transformed, mean-centred, and Pareto-scaled. To ensure robust downstream supervised modelling, we first performed Principal Component Analysis (PCA), Uniform Manifold Approximation and Projection (UMAP), and Non-metric Multidimensional Scaling (NMDS; Euclidean distance) (Supplementary Fig. S1). These unsupervised methods were applied to visualize global data structure, identify outliers, and evaluate treatment-related patterns, and reduce overfitting risk during Orthogonal Partial Least Squares-Discriminant Analysis (OPLS-DA). OPLS-DA was then used to assess metabolic discrimination between treatments in root and leaf tissues separately. Model quality was evaluated using R^2^X, R^2^Y, and Q^2^, with validation by 200 permutation tests. Model significance was assessed by CV-ANOVA at *P*-value < 0.05. Discriminant features were first defined using VIP > 1. High-confidence differential features were defined using VIP > 1, |log_2_FC| > 0.585 (i.e., 1.5-fold difference between treatments), and FDR-adjusted *P* < 0.05 (Welch’s t-test with Benjamini-Hochberg correction). For individual metabolite comparisons, data were log_2_-transformed to improve normality (assessed by the Shapiro-Wilk test) and analysed using Welch’s one-way ANOVA (unequal variances), followed by Games-Howell post hoc tests (Perez-Fons *et al*., 2023); *P* values were adjusted using the Benjamini-Hochberg procedure (FDR), with FDR-adjusted *P* < 0.05 considered significant. To assess functional pathway activity, a Module Activity Score was calculated by averaging the Z-score normalised abundances of annotated metabolites mapped to a specific KEGG module or functional class. Significant differences in module scores were determined using t-tests with FDR correction. Hierarchical clustering (Euclidean distance) was applied to Z-score normalised differential features. Pearson correlation networks were constructed within each tissue across all samples (roots: n = 16; leaves: n = 15) using the high-confidence differential feature set as nodes and analysed using the *igraph* package (Csárdi & Nepusz, 2006) in R. Edges represented significant Pearson correlations among features (|r| > 0.7, *P* < 0.05), and treatment-specific responses were overlaid as node attributes (direction of change relative to control and primary responder, defined as the treatment with the largest absolute log_2_FC). Network modules were identified via the Louvain algorithm (Blondel *et al*., 2008; Rahiminejad *et al*., 2019) and assigned to functional categories based on MSCC classifications and available structural annotations. Network visualisation employed the Fruchterman-Reingold force-directed layout algorithm (Fruchterman & Reingold, 1991). To assess whether VOC chemical classes predict metabolome variation, generalised linear models (GLMs) were fitted to link tissue-specific metabolome PC1 scores with VOC data from *Zhu et al. (2025, preprint)*, including aggregated VOC abundances by chemical class and categorical predictors (dominant sesquiterpene/monoterpene class, treatment). Models were evaluated using both logistic regression (binomial family, χ^2^ test) for binarized outcomes and linear regression (F-test) for continuous outcomes. Raw metabolomics data, processed feature matrices, and custom R scripts are available in the OSF (Open Science Framework) repository (DOI:10.17605/OSF.IO/7Y4E8).

## Results

### Fungal VOCs induce distinct metabolic profiles in poplar roots and leaves

To investigate whether fungal VOCs can induce metabolic reprogramming in the absence of physical contact, we employed a non-contact pot-in-pot system (Fig. 1a) to expose *Populus* × *canescens* roots to VOCs emitted by fungi with different ecological lifestyles: the pathogen *Heterobasidion annosum*, the ectomycorrhizal fungus *Laccaria bicolor*, and the saprotroph *Postia placenta*. Untargeted metabolomics detected 949 metabolic features across all the samples (OSF repository), of which 94 were putatively annotated to chemical structures by MS/MS spectral matching. An additional 263 compounds were annotated based on MS^1^ data, yielding an overall annotation rate of 37.6% (357/949). Unannotated features were classified into functional categories (lipids, secondary metabolites, peptides/amino acids, carbohydrates, and unknowns) using the MSCC method.

**Figure 1.**
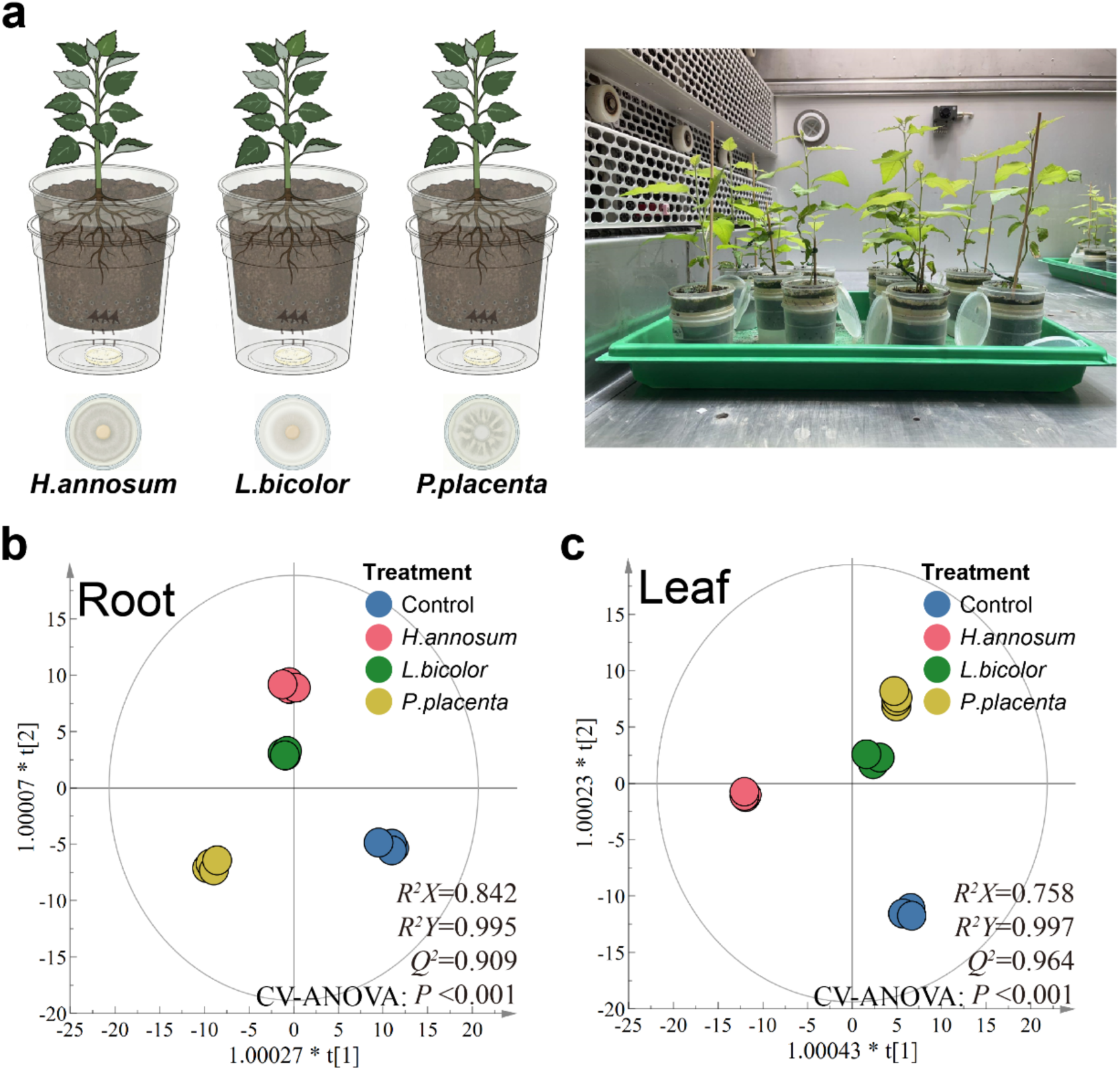
Experimental design and metabolic discrimination of poplar tissues following root exposure to fungal volatile organic compounds (VOCs). (a) Schematic illustration (left) and photograph (right) of the pot-in-pot (PiP) exposure system. *Populus* × *canescens* plants were grown in upper pots with perforated bases and positioned above lower pots containing fungal cultures on MMN medium. Three fungi representing distinct ecological lifestyles were used: the pathogen *H. annosum*, the ectomycorrhizal fungus *L. bicolor*, and the saprotroph *P. placenta*. This setup enabled root exposure to fungal VOCs without physical contact (VOC exposure: n = 6 plants per treatment). (b, c) orthogonal partial least squares–discriminant analysis (OPLS-DA) score plots of root (b) and leaf (c) LC-MS feature profiles after six weeks of VOC exposure. Each point represents one biological replicate included in the metabolomics dataset (n = 4 per treatment; leaf control n = 3). Ellipses indicate the 95% Hotelling’s T^2^ confidence ellipse. Model quality parameters (R^2^X, R^2^Y, Q^2^) and cross-validated ANOVA (CV-ANOVA) significance are shown.

OPLS-DA revealed significant separation in root metabolic profiles among all four groups (R^2^X = 0.842, R^2^Y = 0.995, Q^2^ = 0.909; CV-ANOVA: *P* < 0.001; Fig. 1b). The metabolome of *H. annosum*-exposed roots was very different from controls, while *L. bicolor*- and *P. placenta*-exposed roots differ to a lesser extent, suggesting that pathogen-derived volatiles trigger qualitatively different responses in root tissue than non-pathogenic fungi do. In leaves, the metabolome profiles were similarly discriminated among treatments (R^2^X = 0.758, R^2^Y = 0.997, Q^2^ = 0.964; CV-ANOVA: *P* < 0.001; Fig. 1c). Leaf tissues revealed clear separation among treatments, despite VOC exposure occurring only belowground, indicating systemic metabolic reprogramming. Comparison of Fig. 1b and 1c revealed that *H. annosum* induced the greatest separation from controls in both tissues, whereas the relative positioning of *L. bicolor* and *P. placenta* differed between roots and leaves, suggesting tissue-specific configurations in response to non-pathogenic fungal volatiles.

Principal component analysis (PCA) revealed treatment-associated clustering patterns in roots and leaves (Fig. S1a, b). This organisation was supported by UMAP analyses, which preserve local neighbourhood structure, and by NMDS ordination based on the Euclidean distance of scaled data (Fig. S1c–f). Leaves showed clearer separation between treatments than roots in these ordinations, indicating more coordinated systemic responses, despite the involvement of fewer features and VOC exposure being restricted to the belowground compartment. These convergent patterns across PCA, UMAP and NMDS provided an internal robustness check prior to supervised discrimination and downstream mass feature selection.

### Discriminant feature profiles reveal shared and treatment-associated responses

Based on the OPLS-DA model, mass features with VIP values greater than 1 were selected to examine treatment-related response patterns, yielding 172 mass features in roots and 113 in leaves. Heatmap analysis revealed tissue-dependent metabolic changes (Fig. S2a, d), with compound classes including carbohydrates, lipids, secondary metabolites, peptides, and amino acids. In roots, all fungal treatments clustered together but showed distinct regulation patterns. In leaves, *H. annosum* showed a unique profile dominated by downregulated mass features. Venn diagrams showed that many mass features were shared across all fungal treatments (21 upregulated and 54 downregulated in roots; Fig. S2b, c, e, f), suggesting a common metabolic response to fungal VOCs. Volcano plots further supported the *H. annosum* suppression pattern, with a clear leftward shift in both tissues (Fig. S3).

To focus on statistically robust signals, stricter criteria were applied (VIP > 1, |log_2_FC| > 0.585, FDR-adjusted *P* < 0.05), resulting in 58 and 63 high-confidence mass features in roots and leaves, respectively (Fig. 2). Under these criteria, the shared response was reduced (from 21 to 8 upregulated and from 54 to 6 downregulated in roots; Fig. 2b, c), indicating that many shared mass features had moderate fold changes or high variability. However, treatment-specific patterns remained clear. *H. annosum* uniquely suppressed 21 leaf mass features, over one-third of all high-confidence leaf mass features but showed no unique upregulation in leaves. *P. placenta* retained 11 uniquely upregulated root mass features, showing a distinct activation pattern. *L. bicolor* showed the largest reduction in unique mass features (from 16 to 2 upregulated in roots), suggesting its response overlaps more with other treatments.

**Figure 2.**
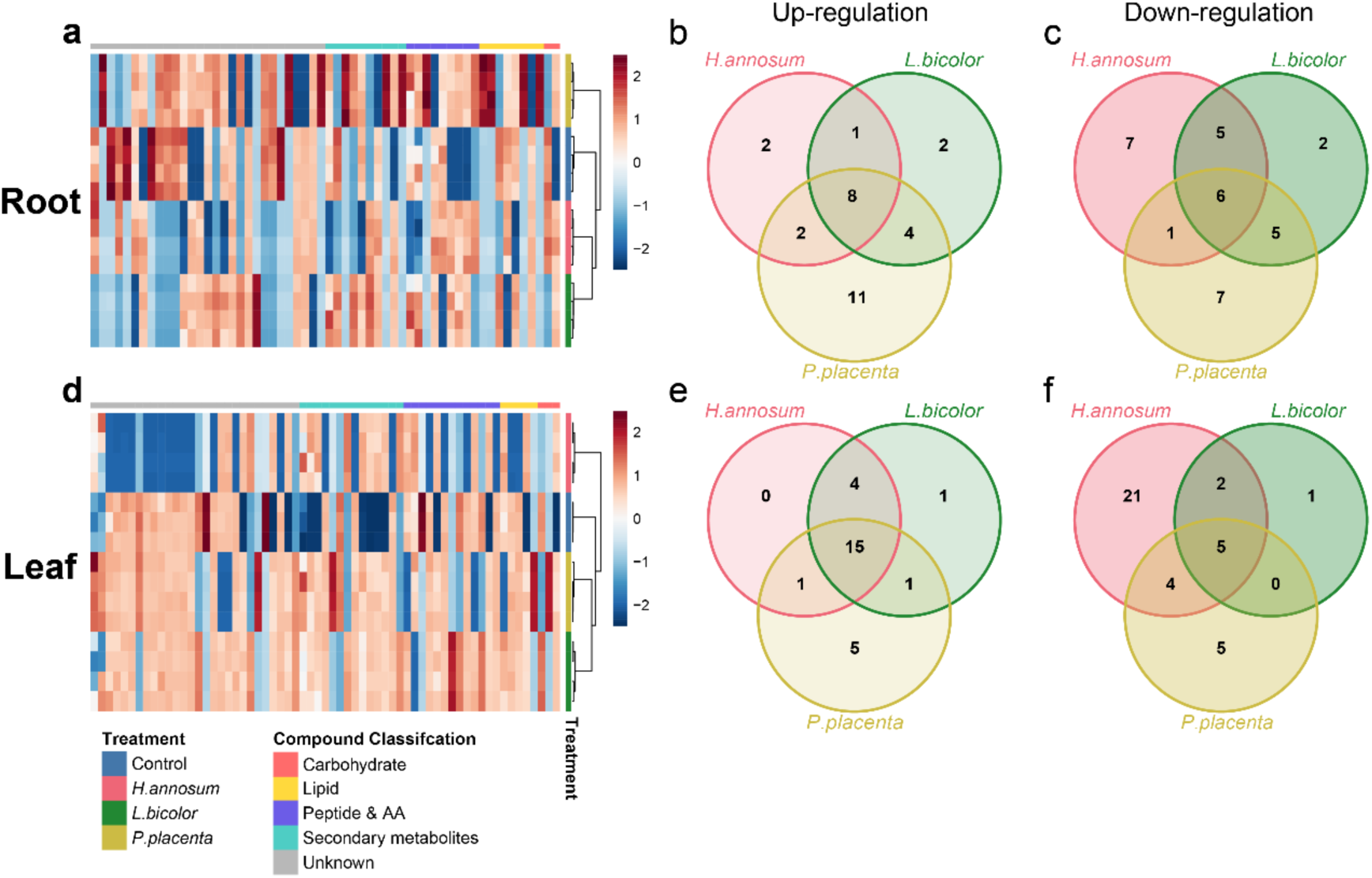
High-confidence differential feature signatures in poplar tissues following root exposure to fungal VOCs. (a, d) Heatmaps showing Z-score normalised abundance of differential features (VIP > 1, |log_2_FC| > 0.585, FDR-adjusted *P* < 0.05) in root (a) and leaf (d) tissues. Hierarchical clustering (Euclidean distance, complete linkage) was applied to both samples (columns) and features (rows). Colour scale indicates relative abundance (red: elevated; blue: reduced). Top annotation bars indicate treatment groups. Left annotation bars show compound classifications based on Multidimensional Stoichiometric Compound Classification (MSCC). (b, c, e, f) Venn diagrams displaying the overlap of upregulated (b, e) and downregulated (c, f) features among treatments in root (b, c) and leaf (e, f) tissues.

### Correlation networks reveal tissue-dependent metabolic co-regulation

To investigate the coordinated regulation of high-confidence differential mass features, we constructed Pearson correlation networks for each tissue and treatment, overlaying treatment-specific responses as node attributes (Fig. 3). These networks exhibited clear structural variations between tissues, revealing distinct metabolic co-regulation architectures that depend on both fungal treatment and tissue type. Leaf metabolic networks consistently exhibited greater connectivity than root networks across all treatments (Fig. 3b, d, f vs. Fig. 3a, c, e), indicating that leaves exhibit more coordinated responses. All three leaf metabolic networks featured modules enriched in secondary metabolites and lipids, suggesting that these classes disproportionately contribute to systemic co-variation patterns.

**Figure 3.**
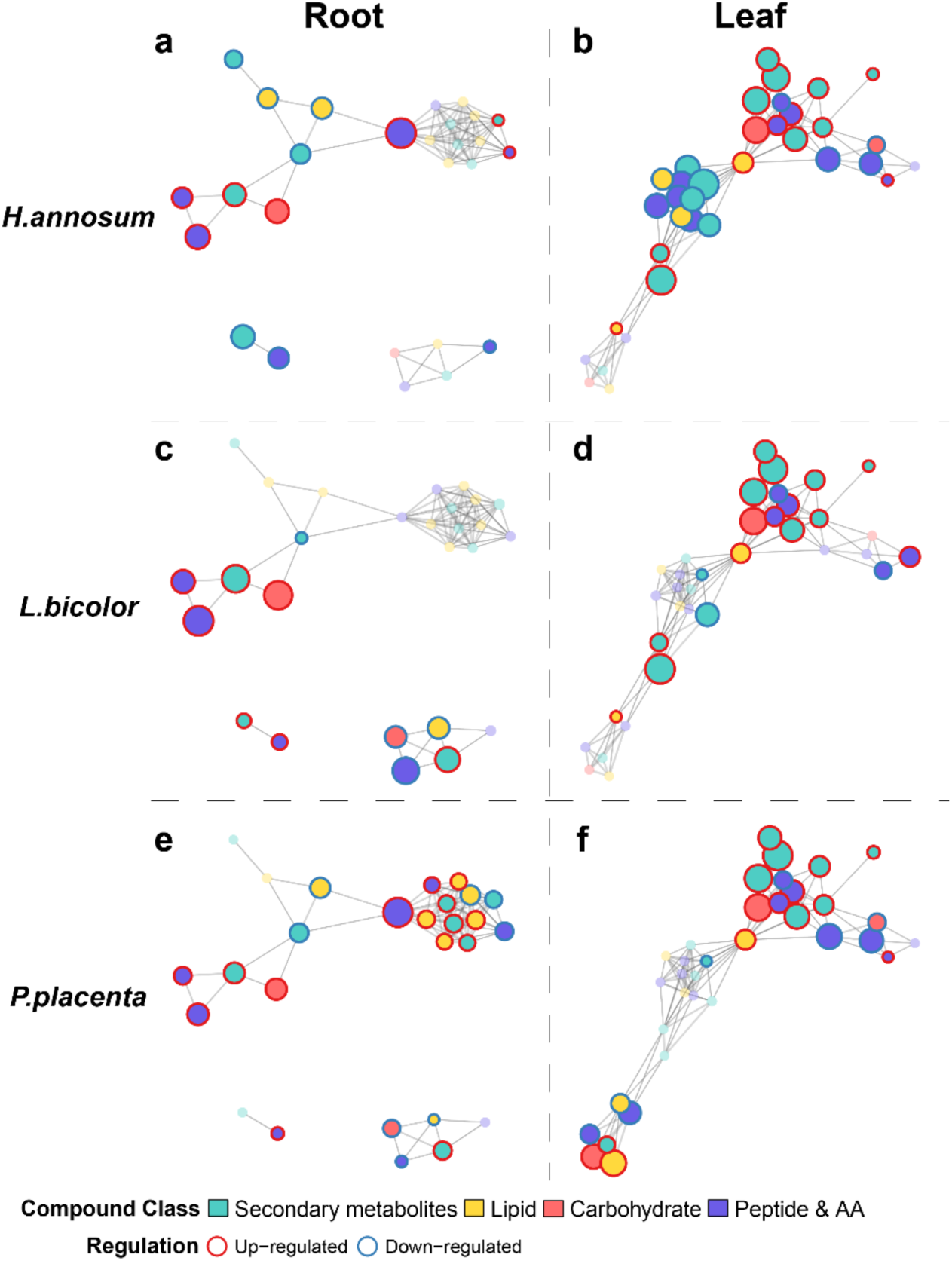
Correlation metabolic networks of high-confidence discriminant mass features reveal tissue-specific metabolic coordination following root exposure to fungal VOCs. Pearson correlation networks were constructed within each tissue across all available samples (roots: n = 16; leaves: n = 15) using the set of high-confidence differential features (variable importance in projection, VIP > 1, |log_2_FC| > 0.585, FDR-adjusted *P* < 0.05) as nodes. Edges indicate strong significant Pearson correlations (|r| > 0.7, *P* < 0.05). Root (a, c, e) and leaf (b, d, f) networks are shown, with treatment-specific overlays for: *H. annosum* (a, b), *L. bicolor* (c, d), and *P. placenta* (e, f). Node fill colours denote compound classes based on Multidimensional Stoichiometric Compound Classification (MSCC). Node border colours indicate the direction of regulation for the corresponding treatment relative to control. Node size is proportional to the absolute log_2_ fold change (|log_2_FC|) for the corresponding treatment comparison. Network layout was generated using the Fruchterman-Reingold algorithm.

Root metabolic networks were sparser overall, but they also revealed treatment-dependent differences. *H. annosum* roots (Fig. 3a) exhibited intermediate connectivity, displaying scattered clusters that primarily contained downregulated mass features. In the *L. bicolor* treatment, the root metabolic network appeared more fragmented (Fig. 3c), with small, isolated clusters consistent with the limited number of unique, high-confidence metabolic changes observed. The *P. placenta* root metabolic network (Fig. 3e) revealed well-defined clusters, including a clear module of jointly up-regulated lipids.

Treatment-associated structures were evident within leaf metabolic networks. The *H. annosum* leaf network (Fig. 3b) contained a large cluster of jointly down-regulated secondary metabolites, reflecting its extensive suppression pattern in the differential feature analysis. In the *L. bicolor* leaf metabolic network (Fig. 3d), treatment-associated structures were weaker overall, with more fragmented connectivity and smaller, dispersed clusters, consistent with the limited number of unique, high-confidence metabolic changes observed. The *P. placenta* leaf network (Fig. 3f) contained multiple modules with mixed regulation, including clusters of upregulated lipids, consistent with its activation-dominated signature.

The distribution of compound classes supported functional grouping: secondary metabolites formed the core of leaf metabolic networks and were often jointly downregulated (particularly in tissues exposed to *H. annosum*), whereas lipids formed distinct modules that were especially prominent in *P. placenta* networks. These architectures suggest that VOC perception is associated with changes at the metabolite level and network-level reorganisation. Treatment-dependent module composition reflects distinct metabolic programmes.

### Representative annotated metabolites reveal tissue-specific functional responses

To clarify the metabolic impact of treatment-associated changes, representative annotated metabolites were visualized using dot plots organised by functional group and KEGG pathway classification (Fig. 4a, b and Table 1). In roots (Fig. 4a), metabolites showed treatment-specific regulation across compound classes: lipids such as ceramides and phosphatidic acids varied among treatments, multiple tripeptides exhibited upregulation that differed by treatment, and phenolic compounds displayed variable profiles across fungal exposures. In leaves (Fig. 4b), salicylic acid 2-O-sulfate (SA-S) showed broad upregulation across all fungal treatments compared to controls. Certain lipids showed *P. placenta*-specific accumulation, while rutin showed contrasting regulation: significantly downregulated by *H. annosum* but maintained or elevated in other treatments.

**Figure 4.**
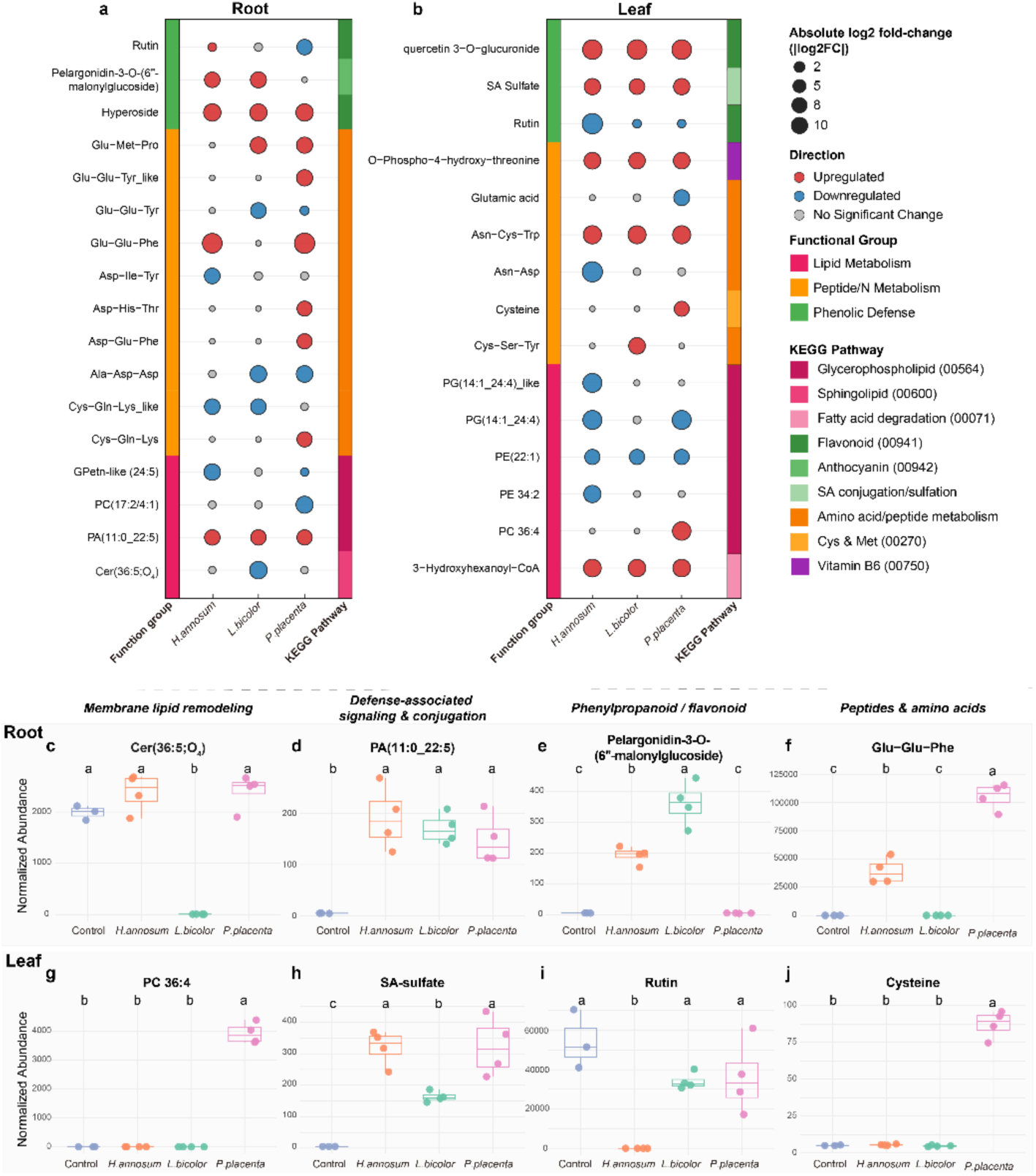
Tissue-specific metabolite responses to fungal volatile organic compounds (VOCs) in poplar roots and leaves. (a, b) Dot plots summarizing responses of selected metabolites in roots (a) and leaves (b) after contact-free root exposure to VOCs emitted by *H. annosum* (pathogen), *L. bicolor* (mutualist), and *P. placenta* (saprotroph). Dot size represents the absolute log_2_ fold-change (|log_2_FC|) relative to controls. Red and blue dots indicate significant up- and down-regulation, respectively (FDR-adjusted *P* < 0.05); grey dots indicate no significant change. Left colour bars denote functional groups; right colour bars indicate KEGG pathway classifications. (c–j) Box plots of median-normalized abundance for representative metabolites, illustrating four response patterns in roots (c–f) and leaves (g–j): membrane lipid remodeling (c, g), defence-associated signalling and conjugation (d, h), phenylpropanoid/flavonoid (e, i), and peptides and amino acids (f, j). Different letters indicate significant differences among treatments (Welch’s one-way ANOVA followed by Games-Howell post hoc tests, FDR-adjusted *P* < 0.05; n = 4 biological replicates per treatment, except leaf controls n = 3).

**Table 1.**
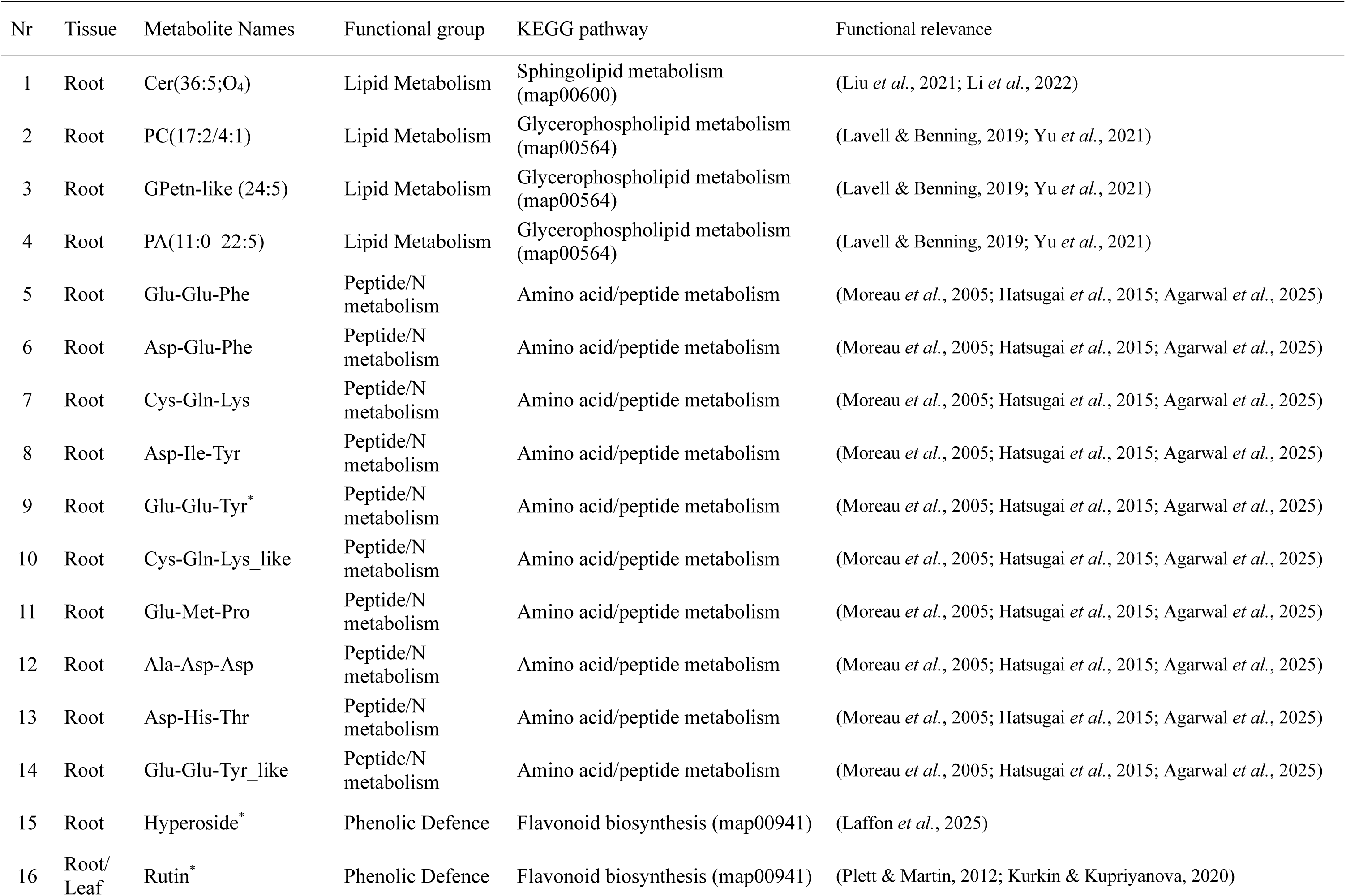
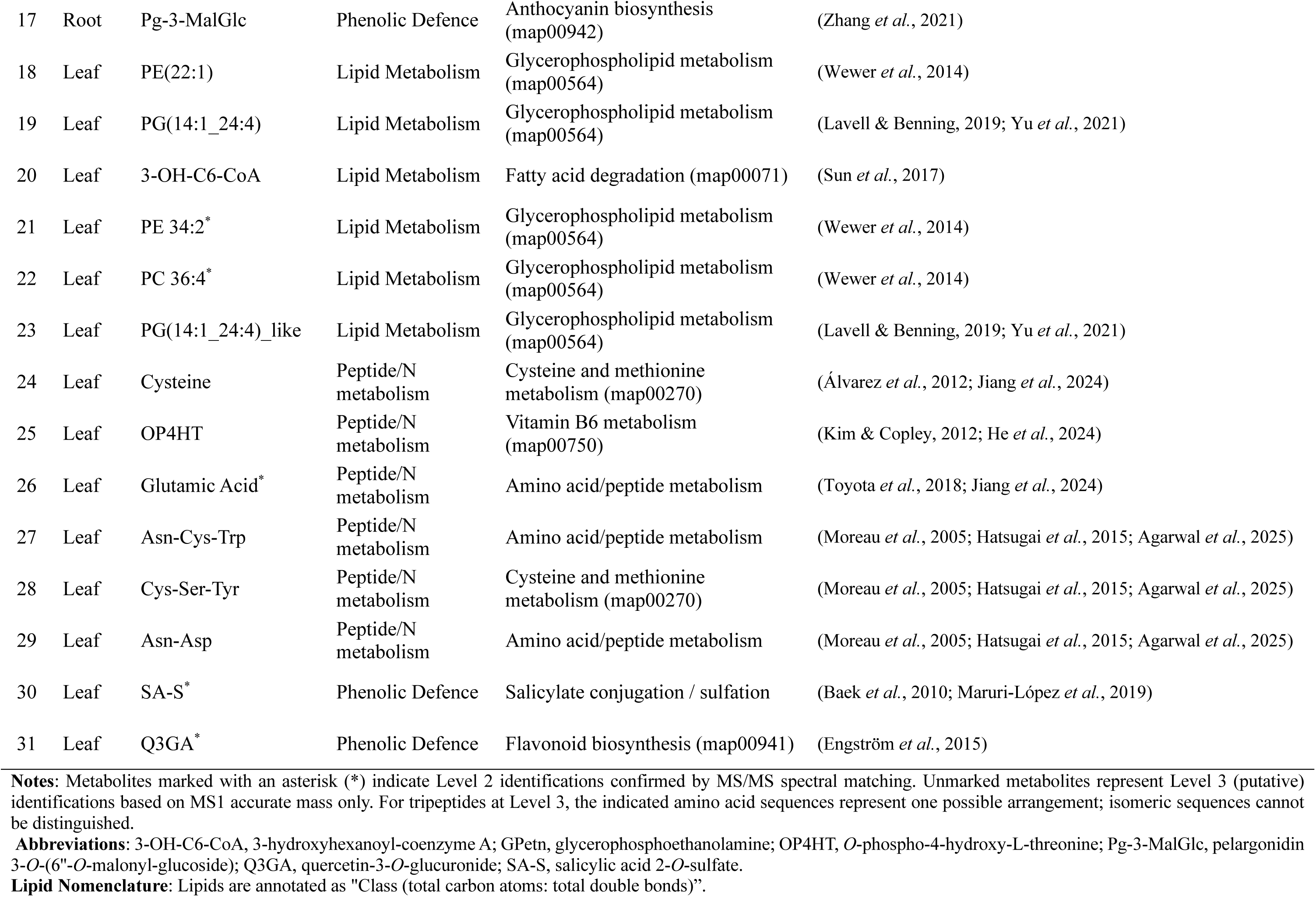
Detailed information of representative differential metabolites in poplar roots and leaves responding to fungal VOCs.

| Nr | Tissue | Metabolite Names | Functional group | KEGG pathway | Functional relevance |
| --- | --- | --- | --- | --- | --- |
| 1 | Root | Cer(36:5;O<sub>4</sub>) | Lipid Metabolism | Sphingolipid metabolism (map00600) | (Liu *et al.*, 2021; Li *et al.*, 2022) |
| 2 | Root | PC(17:2/4:1) | Lipid Metabolism | Glycerophospholipid metabolism (map00564) | (Lavell & Benning, 2019; Yu *et al.*, 2021) |
| 3 | Root | GPetn-like (24:5) | Lipid Metabolism | Glycerophospholipid metabolism (map00564) | (Lavell & Benning, 2019; Yu *et al.*, 2021) |
| 4 | Root | PA(11:0_22:5) | Lipid Metabolism | Glycerophospholipid metabolism (map00564) | (Lavell & Benning, 2019; Yu *et al.*, 2021) |
| 5 | Root | Glu-Glu-Phe | Peptide/N metabolism | Amino acid/peptide metabolism | (Moreau *et al.*, 2005; Hatsugai *et al.*, 2015; Agarwal *et al.*, 2025) |
| 6 | Root | Asp-Glu-Phe | Peptide/N metabolism | Amino acid/peptide metabolism | (Moreau *et al.*, 2005; Hatsugai *et al.*, 2015; Agarwal *et al.*, 2025) |
| 7 | Root | Cys-Gln-Lys | Peptide/N metabolism | Amino acid/peptide metabolism | (Moreau *et al.*, 2005; Hatsugai *et al.*, 2015; Agarwal *et al.*, 2025) |
| 8 | Root | Asp-Ile-Tyr | Peptide/N metabolism | Amino acid/peptide metabolism | (Moreau *et al.*, 2005; Hatsugai *et al.*, 2015; Agarwal *et al.*, 2025) |
| 9 | Root | Glu-Glu-Tyr* | Peptide/N metabolism | Amino acid/peptide metabolism | (Moreau *et al.*, 2005; Hatsugai *et al.*, 2015; Agarwal *et al.*, 2025) |
| 10 | Root | Cys-Gln-Lys_like | Peptide/N metabolism | Amino acid/peptide metabolism | (Moreau *et al.*, 2005; Hatsugai *et al.*, 2015; Agarwal *et al.*, 2025) |
| 11 | Root | Glu-Met-Pro | Peptide/N metabolism | Amino acid/peptide metabolism | (Moreau *et al.*, 2005; Hatsugai *et al.*, 2015; Agarwal *et al.*, 2025) |
| 12 | Root | Ala-Asp-Asp | Peptide/N metabolism | Amino acid/peptide metabolism | (Moreau *et al.*, 2005; Hatsugai *et al.*, 2015; Agarwal *et al.*, 2025) |
| 13 | Root | Asp-His-Thr | Peptide/N metabolism | Amino acid/peptide metabolism | (Moreau *et al.*, 2005; Hatsugai *et al.*, 2015; Agarwal *et al.*, 2025) |
| 14 | Root | Glu-Glu-Tyr_like | Peptide/N metabolism | Amino acid/peptide metabolism | (Moreau *et al.*, 2005; Hatsugai *et al.*, 2015; Agarwal *et al.*, 2025) |
| 15 | Root | Hyperoside* | Phenolic Defence | Flavonoid biosynthesis (map00941) | (Laffon *et al.*, 2025) |
| 16 | Root/<br>Leaf | Rutin* | Phenolic Defence | Flavonoid biosynthesis (map00941) | (Plett & Martin, 2012; Kurkin & Kupriyanova, 2020) |
| 17 | Root | Pg-3-MalGlc | Phenolic Defence | Anthocyanin biosynthesis (map00942) | (Zhang *et al.*, 2021) |
| 18 | Leaf | PE(22:1) | Lipid Metabolism | Glycerophospholipid metabolism (map00564) | (Wewer *et al.*, 2014) |
| 19 | Leaf | PG(14:1_24:4) | Lipid Metabolism | Glycerophospholipid metabolism (map00564) | (Lavell & Benning, 2019; Yu *et al.*, 2021) |
| 20 | Leaf | 3-OH-C6-CoA | Lipid Metabolism | Fatty acid degradation (map00071) | (Sun *et al.*, 2017) |
| 21 | Leaf | PE 34:2* | Lipid Metabolism | Glycerophospholipid metabolism (map00564) | (Wewer *et al.*, 2014) |
| 22 | Leaf | PC 36:4* | Lipid Metabolism | Glycerophospholipid metabolism (map00564) | (Wewer *et al.*, 2014) |
| 23 | Leaf | PG(14:1_24:4)_like | Lipid Metabolism | Glycerophospholipid metabolism (map00564) | (Lavell & Benning, 2019; Yu *et al.*, 2021) |
| 24 | Leaf | Cysteine | Peptide/N metabolism | Cysteine and methionine metabolism (map00270) | (Álvarez *et al.*, 2012; Jiang *et al.*, 2024) |
| 25 | Leaf | OP4HT | Peptide/N metabolism | Vitamin B6 metabolism (map00750) | (Kim & Copley, 2012; He *et al.*, 2024) |
| 26 | Leaf | Glutamic Acid* | Peptide/N metabolism | Amino acid/peptide metabolism | (Toyota *et al.*, 2018; Jiang *et al.*, 2024) |
| 27 | Leaf | Asn-Cys-Trp | Peptide/N metabolism | Amino acid/peptide metabolism | (Moreau *et al.*, 2005; Hatsugai *et al.*, 2015; Agarwal *et al.*, 2025) |
| 28 | Leaf | Cys-Ser-Tyr | Peptide/N metabolism | Cysteine and methionine metabolism (map00270) | (Moreau *et al.*, 2005; Hatsugai *et al.*, 2015; Agarwal *et al.*, 2025) |
| 29 | Leaf | Asn-Asp | Peptide/N metabolism | Amino acid/peptide metabolism | (Moreau *et al.*, 2005; Hatsugai *et al.*, 2015; Agarwal *et al.*, 2025) |
| 30 | Leaf | SA-S* | Phenolic Defence | Salicylate conjugation / sulfation | (Baek *et al.*, 2010; Maruri-López *et al.*, 2019) |
| 31 | Leaf | Q3GA* | Phenolic Defence | Flavonoid biosynthesis (map00941) | (Engström *et al.*, 2015) |

**Notes:** Metabolites marked with an asterisk (\*) indicate Level 2 identifications confirmed by MS/MS spectral matching. Unmarked metabolites represent Level 3 (putative) identifications based on MS1 accurate mass only. For tripeptides at Level 3, the indicated amino acid sequences represent one possible arrangement; isomeric sequences cannot be distinguished.
**Abbreviations:** 3-OH-C6-CoA, 3-hydroxyhexanoyl-coenzyme A; GPetn, glycerophosphoethanolamine; OP4HT, *O*-phospho-4-hydroxy-L-threonine; Pg-3-MalGlc, pelargonidin 3-*O*-(6"-*O*-malonyl-glucoside); Q3GA, quercetin-3-*O*-glucuronide; SA-S, salicylic acid 2-*O*-sulfate.
**Lipid Nomenclature:** Lipids are annotated as "Class (total carbon atoms: total double bonds)".

Box plot analysis (Fig. 4c–j; adj. *P* < 0.05) confirmed distinct response patterns across functional groups. For membrane lipid remodelling, root Cer(36:5;O4) was significantly lower in *L. bicolor*-exposed plants than in the other three groups (Fig. 4c), while leaf PC(36:4) was significantly higher only under *P. placenta* treatment (Fig. 4g). For defence-associated signalling and conjugation, root PA(11:0_22:5) was significantly elevated in all three fungal treatments compared to controls (Fig. 4d), while leaf SA-S similarly showed significant elevation across all fungal treatments (Fig. 4h). For phenylpropanoid/flavonoid metabolites, root pelargonidin-3-O-(6″-malonylglucoside) levels were highest in *L. bicolor*-exposed plants and were also significantly elevated in *H. annosum*-exposed plants compared to controls and *P. placenta* (Fig. 4e), while leaf rutin was significantly suppressed only in *H. annosum*-exposed plants (Fig. 4i). For peptides and amino acids, root Glu-Glu-Phe levels were highest in *P. placenta*-exposed plants, with *H. annosum* also significantly higher than controls and *L. bicolor* (Fig. 4f), and leaf cysteine was significantly higher only in *P. placenta* (Fig. 4j). Complete response profiles grouped by treatment specificity are provided in Fig. S5.

## Discussion

### Volatile-mediated recognition drives systemic reprogramming

Fungal VOCs can induce systemic, lifestyle-associated physiological reprogramming in *Populus × canescens* even in the absence of physical contact. The metabolomics analysis presented here extends our earlier volatile profiling work (Zhu et al., 2025, preprint) by examining how lifestyle-structured fungal VOC signatures translate into downstream metabolic changes in plant tissues. Using the same pot-in-pot exposure system and plant material, we characterized the root and leaf metabolome after six weeks of VOC exposure, providing a complementary view of plant responses at the molecular level. Across fungi representing pathogenic, saprotrophic, and mutualistic lifestyles, poplars translated airborne chemical cues into distinct and persistent metabolic states. This supports the hypothesis that volatile-mediated surveillance shapes whole-plant physiology prior to contact (Rosenkranz *et al*., 2021). Specifically, responses comprised a conserved core signature across all fungal VOC exposures together with three fungus-associated systemic modes: suppression-dominated *(H. annosum*), lipid-centred activation (*P. placenta*), and restrained compatibility-oriented reprogramming (*L. bicolor*) (Fig. 5).

**Figure 5.**
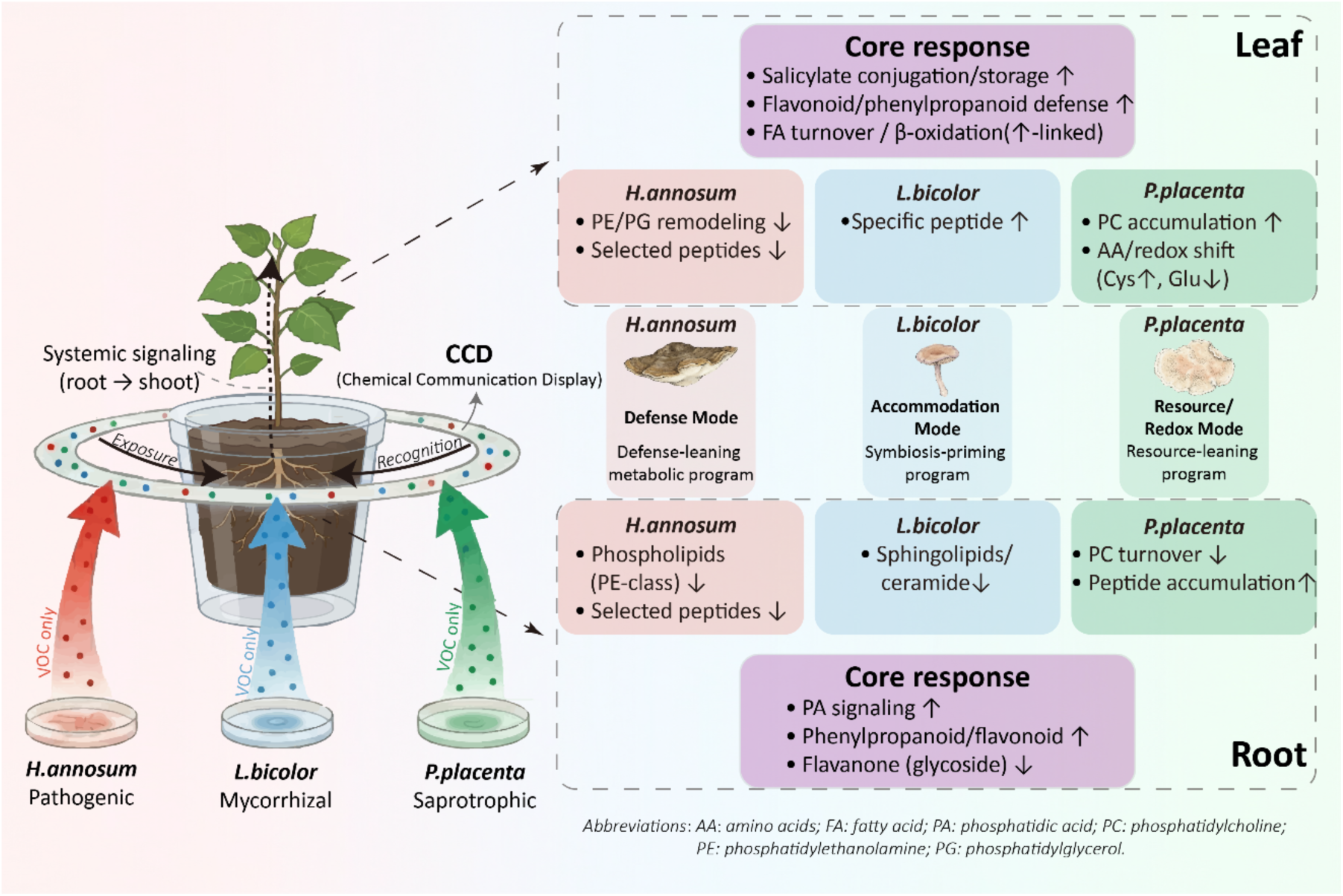
Conceptual model of systemic metabolic reprogramming in poplar roots and leaves triggered by fungal VOCs. The diagram summarizes tissue-specific responses into three ecological modes: Defence Mode (*H. annosum*), Accommodation Mode (*L. bicolor*), and Resource/Redox Mode (*P. placenta*). The upper panel summarizes leaf responses, and the lower panel summarizes root responses. Purple boxes highlight “Core responses” conserved across all treatments, while coloured boxes detail specific metabolic signatures. Dashed arrows from root to leaf indicate systemic root-to-shoot signalling. Upward (↑) and downward (↓) arrows denote regulation direction relative to controls.

### Chemical communication displays (CCDs) provide recognition cues

Volatile-mediated discrimination can be interpreted as an early recognition layer that decodes chemical communication displays (CCDs) (Fig. 5). In the paired volatilome dataset, below-ground headspace profiles remained stable and lifestyle-associated over time, whereas above-ground volatile profiles exhibited greater diversity and partial convergence (Zhu *et al*., 2025, preprint). Our metabolomics results extend this observation by showing that sustained exposure to these airborne blends translates into coordinated systemic metabolic states, comprising a conserved core response alongside lifestyle-associated modes across roots and leaves (Fig. 5).

CCD theory emphasizes that biological meaning can reside in blend composition, ratios, and covariation, rather than in single ’marker’ compounds (Junker *et al*., 2018). GLM analyses linking VOC profiles (from Zhu *et al.,* 2025, preprint) to metabolome PC1 scores supported this interpretation (see Table S1): no individual VOC chemical class consistently predicted metabolome variation, whereas sesquiterpene class composition significantly predicted leaf response (χ^2^ = 10.68, *P* = 0.014) and treatment identity predicted both root (χ^2^ = 11.09, *P* = 0.011) and leaf (F = 4.84, *P* = 0.022) metabolome variation. These results suggest that plants decode lifestyle-associated cues from blend-level VOC architecture rather than individual compounds. Consistent with this, Guo *et al*. (2021) showed that fungal VOC patterns can predict trophic mode and lifestyle across taxa, and quantified phenotypic integration (i.e. compound covariation) within volatilomes. From this perspective, abundant compound classes such as sesquiterpenes are best understood as contributors to CCD architecture and identity encoding, while hormone-weighted physiological states emerge downstream through systemic integration.

### Species-specific vs. lifestyle-associated responses

Our experimental design included one representative fungal species per ecological lifestyle. The observed metabolic responses are therefore fungus-specific, though the patterns align with expectations based on lifestyle categories. Several lines of evidence support a lifestyle-associated interpretation: (i) the strong lifestyle structuring of fungal VOC profiles within the same experiment (Zhu et al., 2025, preprint), (ii) cross-taxon volatilomics studies showing lifestyle-correlated emission patterns (Guo *et al.,* 2021), and (iii) the functional coherence of the plant responses observed here: suppression by a pathogen, activation by a saprotroph, and restrained responses to a mutualist. While these findings are consistent with lifestyle-mediated recognition, we acknowledge that species identity and lifestyle cannot be fully disentangled within a single-species-per-lifestyle design. Future studies incorporating multiple phylogenetically diverse taxa per lifestyle will further test the generality of these patterns.

### Shared metabolic responses across fungal VOCs

Beyond lifestyle contrasts, Grey poplar exhibited a shared VOC-responsive component, particularly in roots. This suggests that common fungal airborne chemistry can elicit conserved metabolic adjustments. Two metabolites illustrate this core response at the pathway level: phosphatidic acid [PA(11:0_22:5)] accumulated in roots across all three fungal treatments, consistent with its role as an early stress signal linked to MAPK-mediated signalling (Zhou *et al*., 2022) and fungal defence responses (Zhao *et al*., 2013). The accumulation of salicylic acid 2-O-sulfate (SA-S) reflects a broad strategy of salicylate conjugation and mobilization in response to fungal cues, aimed at balancing immune activation (Baek *et al*., 2010; Maruri-López *et al*., 2019). Within the CCD concept, such shared responses are compatible with plants responding to conserved ’common-fungal’ modules in VOC mixtures, while fungal-specific outcomes depend on the context and structure of the mixture.

### Pathogen VOCs induce suppression-dominated reprogramming

Pathogen-derived VOC cues from *H. annosum* induced the strongest deviation from controls and a suppression-dominated metabolic phenotype, particularly in leaves. This net inhibitory outcome following prolonged pre-contact exposure aligns with the suppression of secondary metabolism associated with pathogens, as described for contact-dependent pathogenicity mechanisms (Xu *et al*., 2019; Shang *et al*., 2024). Data on paired volatilome dynamics from the same experiment also support a phased response scenario involving early enrichment of green leaf volatiles (GLVs), followed by later enrichment of methyl salicylate (MeSA) in leaves (Zhu *et al*., 2025, preprint). While MeSA is best interpreted as a signalling proxy rather than a direct measure of free salicylic acid (SA) pools, the convergence of time-resolved VOC readouts with persistent metabolome suppression supports the interpretation that pathogen VOCs bias poplar towards a sustained defensive configuration before contact, rather than eliciting only transient alarm responses.

### Saprotroph VOCs trigger lipid-centred activation

VOC cues derived from *P. placenta* produced an intermediate but distinct outcome characterized by comparatively stronger metabolic activation, with prominent lipid-associated changes and selective shifts in nitrogen and sulfur metabolism. One reasonable ecological interpretation is that VOC mixtures associated with saprotrophs signal an environmental context linked to substrate turnover, oxidative processes, and microbial competition (Boddy & Hiscox, 2016; Mali *et al*., 2019). In response to such cues, plants may adjust membrane lipid composition to maintain cellular integrity and prepare for potential biotic challenges. Phospholipids such as phosphatidic acid (PA) function both as structural membrane components and as signalling molecules that coordinate stress responses (Zhao *et al*., 2013; Zhou *et al*., 2022). The accumulation of PA and other lipid classes observed here suggests that *P. placenta* VOCs trigger preparatory membrane remodelling, potentially priming the plant for competitive or oxidative conditions associated with saprotrophic activity (Laupheimer *et al*., 2023). This interpretation is consistent with the ecological flexibility observed in fungi with dual saprotrophic and biotrophic capacities, where some saprotrophs colonise living roots and some mycorrhizal fungi decompose deadwood (Smith *et al*., 2017; Purahong *et al*., 2025). In the paired volatilome dataset, the abundance and composition of VOC signatures associated with saprotrophs changed across the exposure period (Zhu *et al*., 2025, preprint). While our metabolomics provides an endpoint snapshot, the combination of time-resolved VOC dynamics with lipid-centred metabolic activation is consistent with sustained adjustment rather than acute defence commitment.

### Mutualist VOCs elicit restrained, compatibility-oriented programming

VOC cues derived from the mutualist *L. bicolor* elicited restrained reprogramming consistent with compatibility, with comparatively few unique changes to treatment and substantial overlap with other fungal exposures. This moderation is consistent with programmes of mutualistic interactions that minimize host disruption and rely on immune regulation rather than strong defence escalation (Kloppholz *et al*., 2011; Plett *et al*., 2014; Casarrubia *et al*., 2016). In the same experiment, exposure to *L. bicolor* was associated with low overall VOC emission rates and limited induction of defence-associated leaf volatiles (Zhu *et al*., 2025, preprint). This supports the view that ectomycorrhizal VOC CCDs are not perceived as acute danger signals. Instead, they can stimulate lateral root formation in the putative host plant even before physical contact occurs (Ditengou *et al*., 2015). A notable signal at the pathway level in our dataset was the reduction of the ceramide [Cer(36:5;O_4_)] specific to mutualists in roots, consistent with the dampening of sphingolipid-linked stress signalling. Considering the roles of ceramides in defence and programmed cell death (Liu *et al*., 2021; Li *et al*., 2022), this reduction is consistent with the idea of adjustment oriented towards compatibility and is consistent with evidence that the *Populus*-*L. bicolor* symbiosis involves fine-scale jasmonate tuning via effectors such as MiSSP7 (Plett *et al*., 2014; Daguerre *et al*., 2020).

### Evidence for root-shoot integration and systemic perception

One striking feature of the system is the strength of root-shoot integration despite strict belowground exposure. Despite VOC delivery being restricted to the root zone, leaf metabolomes discriminated treatments, suggesting that belowground CCD decoding is integrated into the metabolic state of the whole plant. Alongside the paired volatilome results, in which belowground VOC patterns remained sharply distinct while leaf VOC profiles partially converged (Zhu *et al*., 2025, preprint), this supports a spatial organisation in which the roots act as the primary chemosensory interface, exposed to persistent, lifestyle-structured VOC blends. In contrast, the leaves integrate these cues alongside photosynthetic and whole-plant demands. Identifying the signalling routes that connect root VOC perception to systemic metabolic outcomes will require time-resolved sampling and targeted readouts of candidate long-distance signals.

### Mechanistic implications and limitations

Mechanistically, responses in the absence of contact imply perception routes beyond classical MAMP (microbe-associated molecular pattern) recognition (Jones & Dangl, 2006). Volatile perception may involve rapid membrane-associated events, including Ca²⁺ influx, which couple early sensing to transcriptional and hormonal adjustments (Zebelo *et al*., 2012; Liu *et al*., 2025). Furthermore, volatile-induced Ca²⁺ signatures have been demonstrated by real-time imaging in Arabidopsis (Aratani *et al*., 2023). The CCD framework further predicts that plants may respond to blend-level properties, ratios and co-varying modules, rather than to individual compounds, providing a plausible route to robust discrimination even when single VOCs vary. However, mechanistic inference is constrained by incomplete metabolite annotation (approximately 37%) and the chronic six-week exposure design, which likely integrates direct VOC-triggered regulation with downstream physiological adjustment. Unsupervised ordinations identified treatment-related patterns before supervised modelling. Supervised feature selection and correlation network reconstruction should be considered as a means of generating hypotheses, with the more conservative basis for interpretation being provided by the stricter FDR-filtered differential features.

## Conclusions: Fungal VOCs shape anticipatory plant metabolism

By pairing volatilomics and metabolomics data from the same contact-free exposure experiment (Zhu *et al*., 2025, preprint), we can establish a direct link between lifestyle-structured fungal VOC cues and systemic metabolic reprogramming in a woody host. Pathogen-associated VOC CCDs bias poplar towards strong, suppression-dominated metabolic shifts, whereas saprotroph-associated CCDs promote lipid-centred metabolic adjustments. In contrast, mutualist-associated CCDs elicit restrained, compatibility-consistent reprogramming, with selective modulation of stress-associated pathways. The results support the idea that trees use belowground VOC chemistry - or fungal CCDs - to anticipate interaction context and adjust systemic metabolism before physical contact occurs.

## Acknowledgements

We would like to thank the EUS staff, especially Peter Kary and Armin Richter, for their technical support in running the climate chamber experiments. We would also like to thank Karin Pritsch (EUS) for supplying the fungal cultures and cultivating the fungi. In this manuscript, AI was used for literature searches (ChatGPT) and linguistic revision (DeepL).

## Author contributions

PZ, MR, and JPS. designed research; PZ performed research with IZ and PBSP, (LC-MS); PZ analysed data; PZ, MR, AG, and JPS interpreted data. PZ wrote the paper with input from all authors.

## Data availability

The raw metabolomics data, all processed feature matrices, and the custom R scripts are publicly available in the OSF (Open Science Framework) repository (DOI:10.17605/OSF.IO/7Y4E8).

## Supporting Information

**Figure S1.**
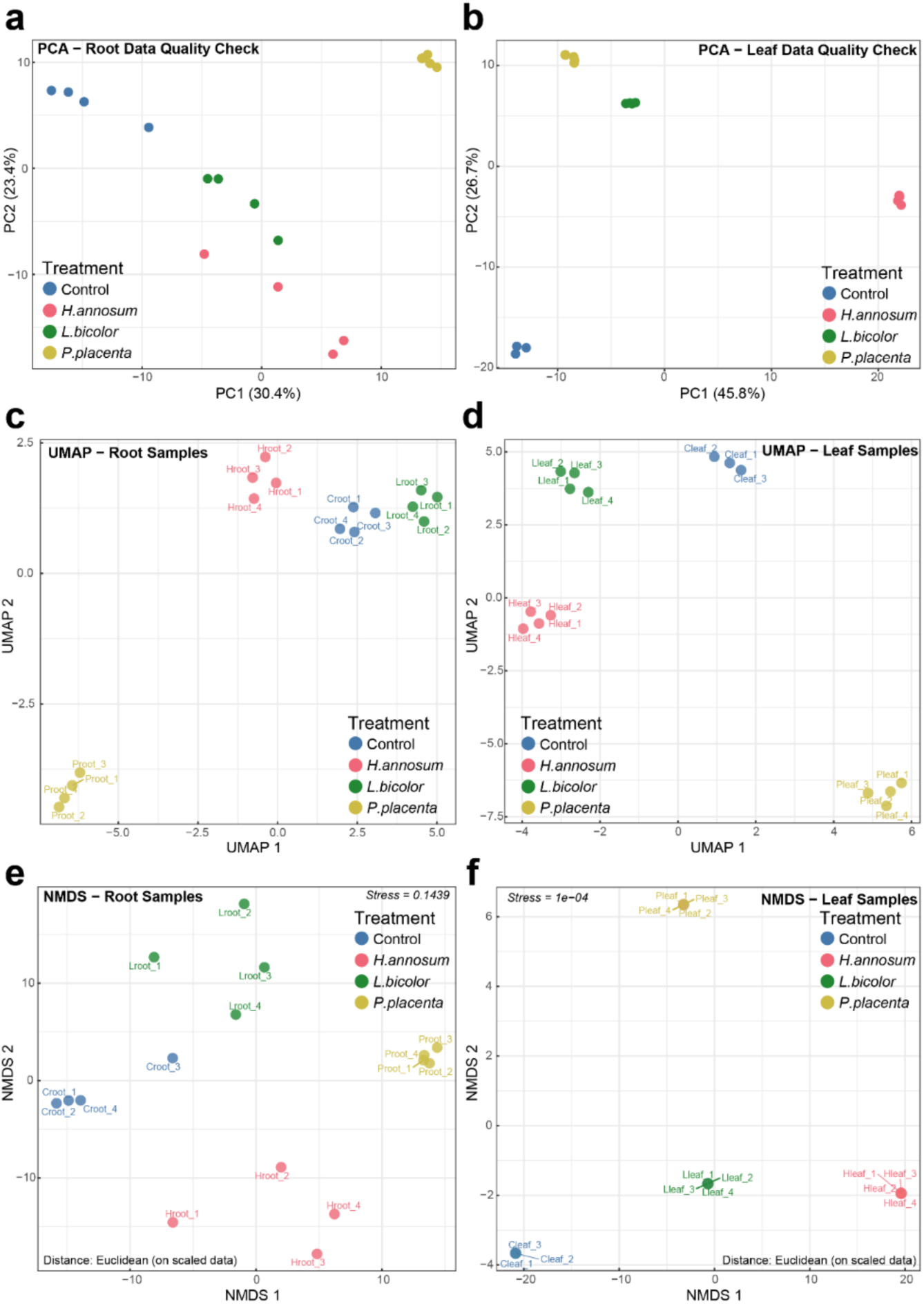
Multivariate visualisation of feature profiles in poplar tissues following root exposure to fungal VOCs. Dimensionality reduction analyses of feature profiles in root (left column) and leaf (right column) tissues using three complementary approaches. (a, b) Principal Component Analysis (PCA) score plots showing sample distribution along PC1 and PC2, with variance explained indicated on each axis. (c, d) Uniform Manifold Approximation and Projection (UMAP) plots preserving local neighbourhood structure. (e, f) Non-metric Multidimensional Scaling (NMDS) ordination based on Euclidean distance of scaled data; stress values are indicated. Colours represent treatment groups: Control (blue), *H. annosum* (pink), *L. bicolor* (green), and *P. placenta* (yellow). Sample labels indicate treatment and replicate number.

**Figure S2.**
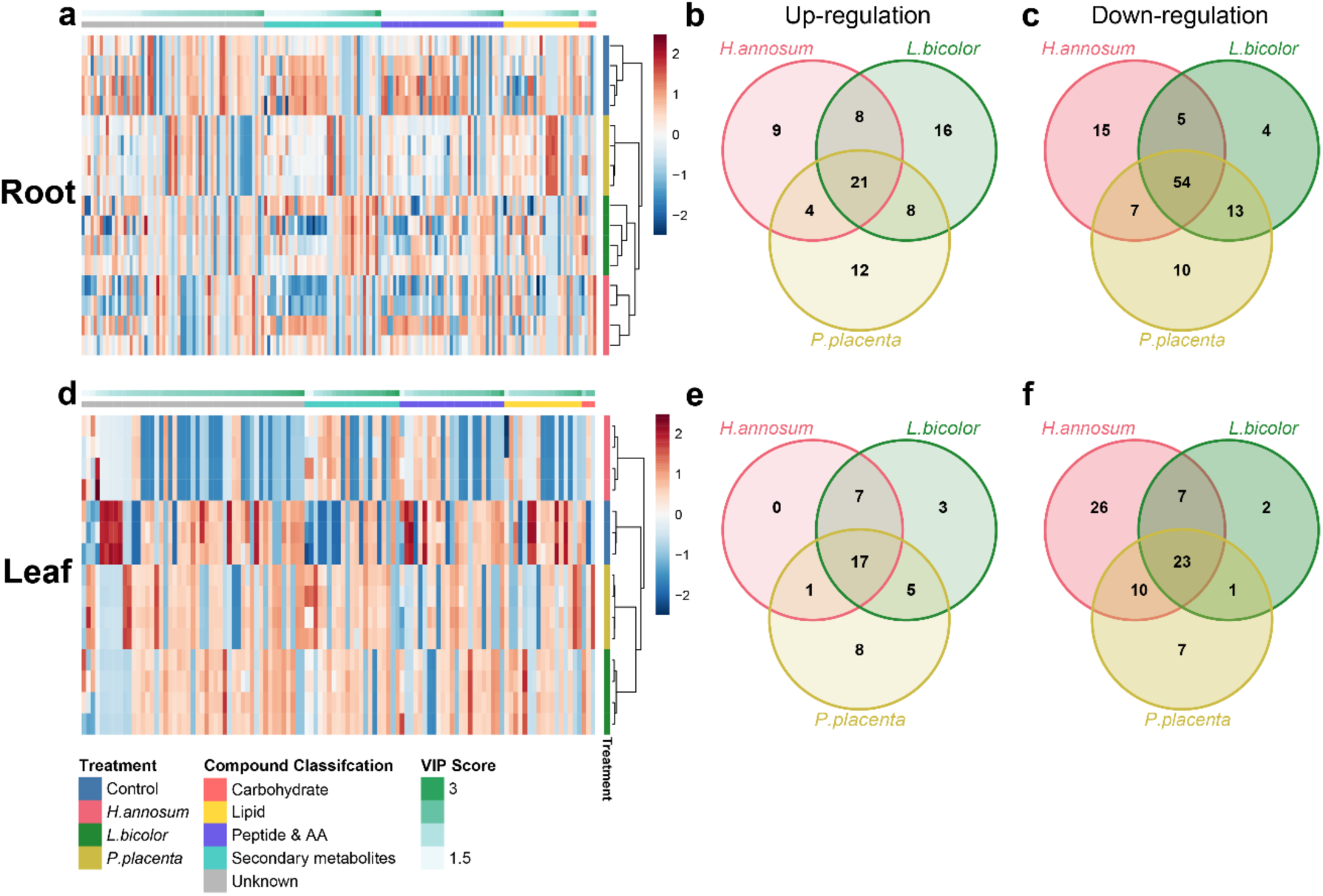
Hierarchical clustering and overlap analysis of discriminant features in poplar tissues following root exposure to fungal VOCs. (a, d) Heatmaps showing Z-score normalised abundance of discriminant features (variable importance in projection, VIP > 1) in root (a) and leaf (d) tissues. Hierarchical clustering (Euclidean distance, complete linkage) was applied to both samples (columns) and features (rows). Colour scale indicates relative abundance (red: elevated; blue: reduced). Top annotation bars indicate treatment groups. Left annotation bars show VIP scores (green gradient) and compound classifications based on Multidimensional Stoichiometric Compound Classification (MSCC). (b, c, e, f) Venn diagrams displaying the overlap of upregulated (log_2_FC > 0.585) (b, e) and downregulated (log_2_FC < -0.585) (c, f) features among treatments in root (b, c) and leaf (e, f) tissues.

**Figure S3.**
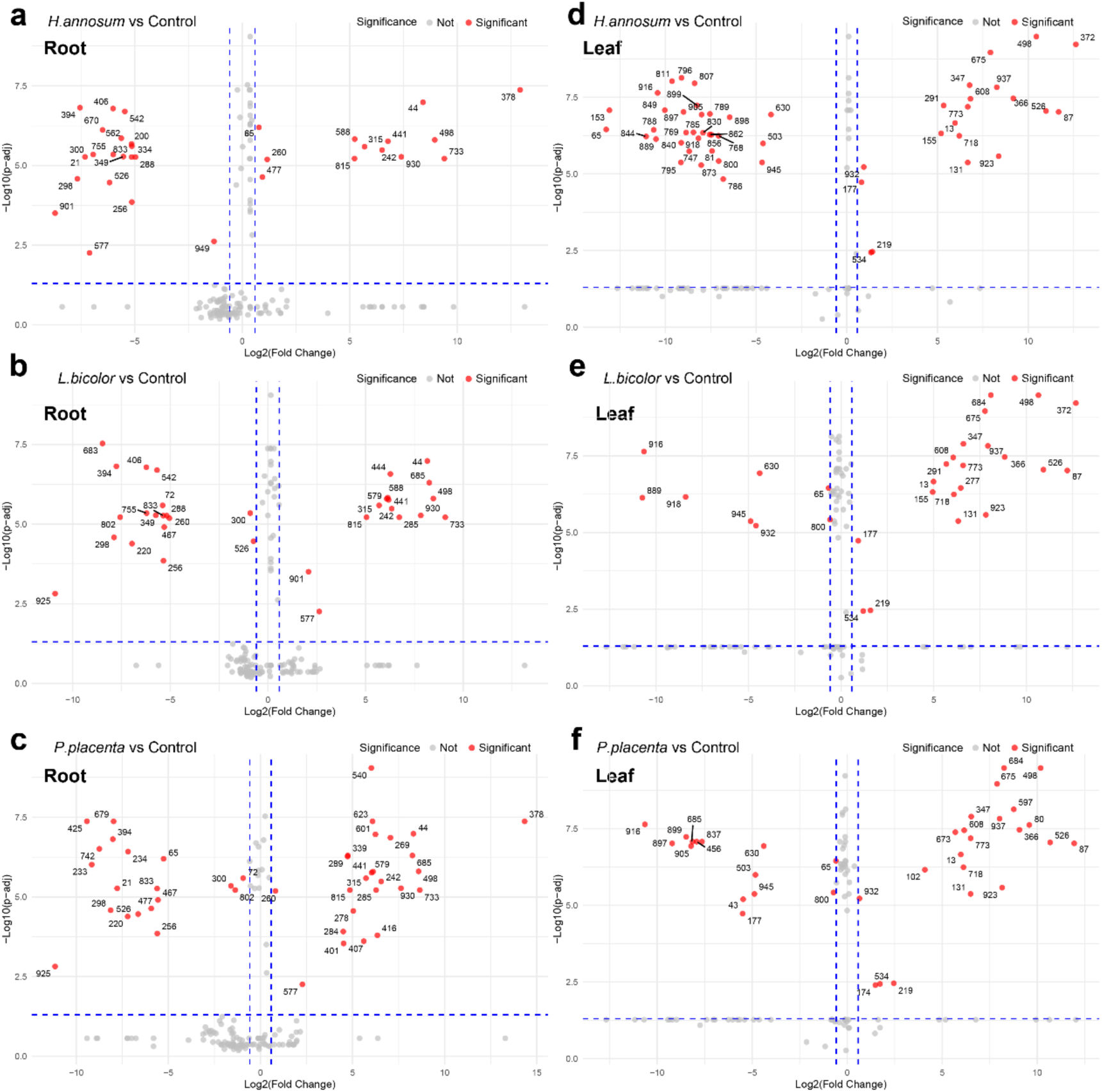
Volcano plots of discriminant features in poplar tissues following root exposure to fungal VOCs. Volcano plots displaying log_2_(fold change) versus – log_10_ (FDR-adjusted *P*-value) for discriminant features (VIP > 1) in root (a-c) and leaf (d-f) tissues. Comparisons are shown for *H. annosum* vs. Control (a, d), *L. bicolor* vs. Control (b, e), and *P. placenta* vs. Control (c, f). Vertical dashed lines indicate fold change thresholds (|log_2_FC| = 0.585); horizontal dashed lines indicate the significance threshold (FDR-adjusted *P* < 0.05). Red points represent differential features meeting both criteria; grey points indicate features not meeting one or both thresholds. Numbers denote feature identifiers.

**Figure S4.**
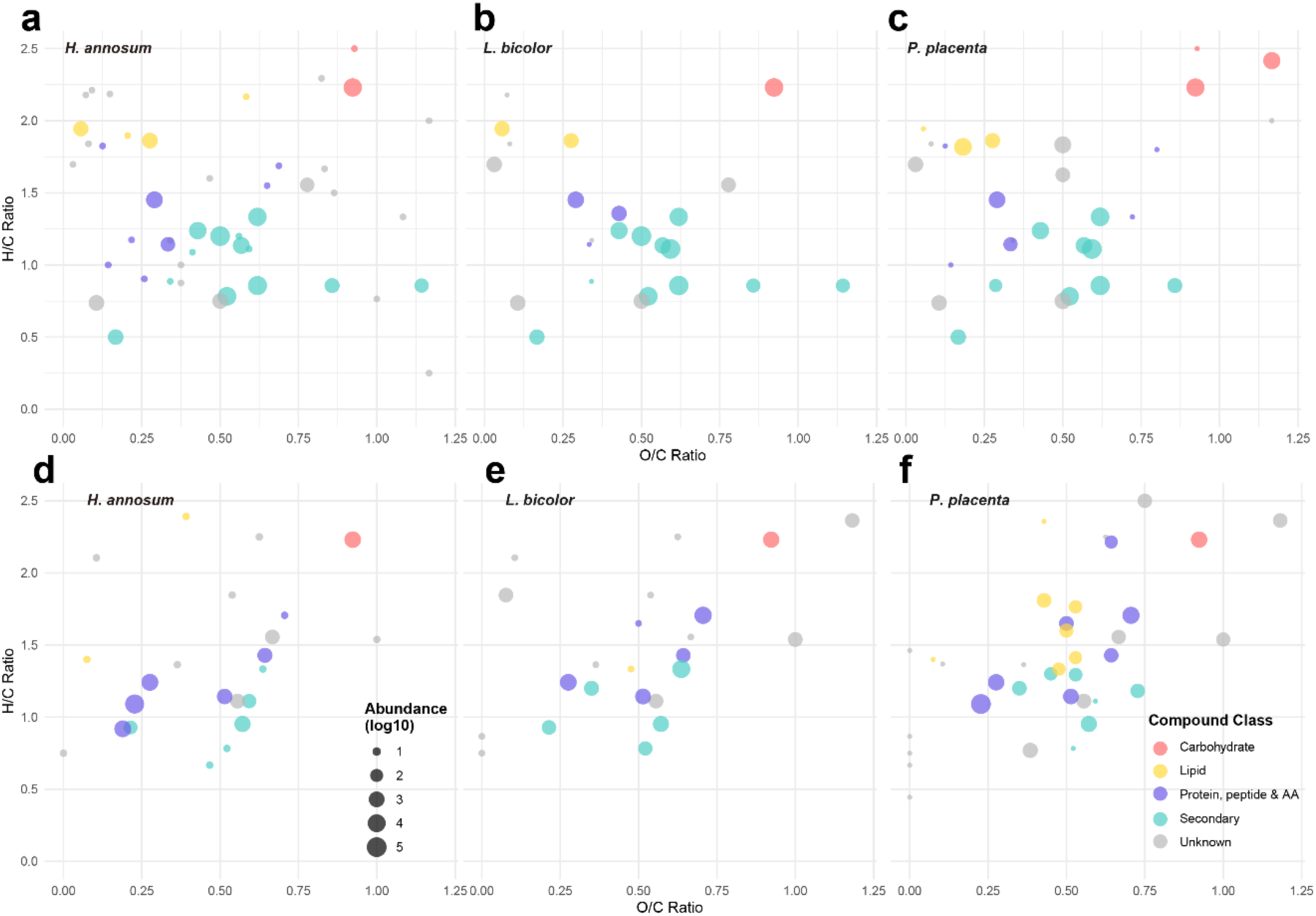
Van Krevelen diagrams of differential features in poplar tissues following root exposure to fungal VOCs. Van Krevelen plots displaying H/C versus O/C elemental ratios of differential features (VIP > 1, |log_2_FC| > 0.585, FDR-adjusted *P* < 0.05) in leaf (a-c) and root (d-f) tissues. Panels show responses to *H. annosum* (a, d), *L. bicolor* (b, e), and *P. placenta* (c, f). Node colours indicate compound classifications based on Multidimensional Stoichiometric Compound Classification (MSCC): carbohydrates (pink), lipids (yellow), proteins/peptides/amino acids (purple), secondary metabolites (cyan), and unknown compounds (grey). Node size reflects log₁₀-transformed feature abundance.

**Figure S5.**
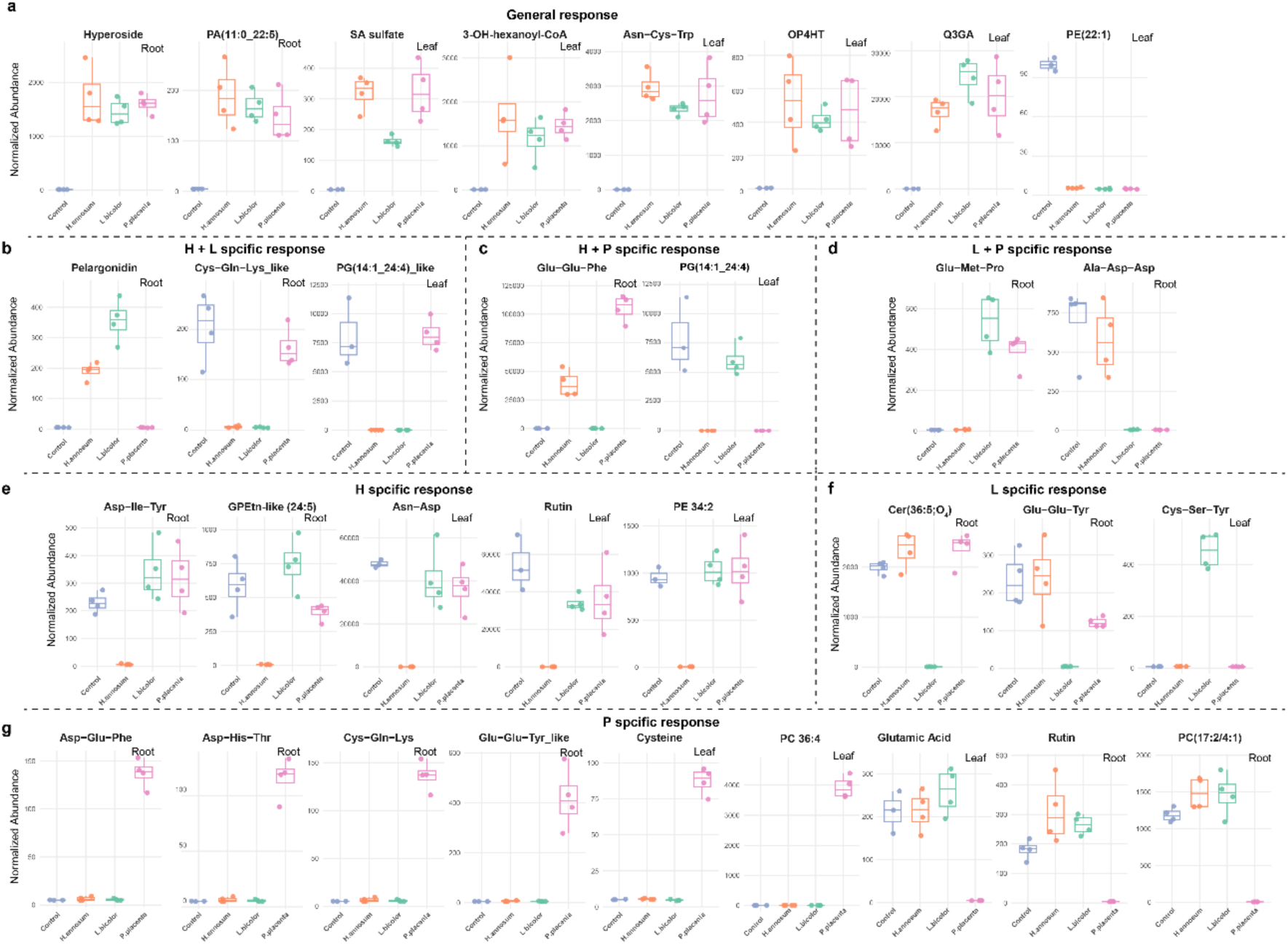
Complete high-confidence feature response profiles grouped by treatment specificity. Boxplots displaying normalised abundance of all differentially regulated features (VIP > 1, |log_2_FC| > 0.585, FDR-adjusted *P* < 0.05) grouped by their response specificity to fungal VOC treatments. H, L, and P represent VOC treatments from *H. annosum*, *L. bicolor*, and *P. placenta*, respectively. (a) General response (regulated by H, L, and P), (b-d) pairwise shared responses (H + L, H + P, L + P), and (e-g) treatment-specific responses (H-only, L-only, P-only). Colours: Control (Purple blue), *H. annosum* (Orange), *L. bicolor* (Green), *P. placenta* (Pink). Tissue type indicated as Root or Leaf. N = 4 biological replicates (n = 3 for leaf control). See Figure 4 for representative examples and summary bubble plots.

**Table S1.** GLM and LM analysis of VOC predictors for plant metabolome variation. VOC data for statistical analysis were derived from Zhu et al. (2025, preprint).

| Term | df | Root PC1_high<br>$\chi^2$ (p) | Leaf PC1_high<br>$\chi^2$ (p) | Root PC1<br>F (p) | Leaf PC1<br>F (p) |
| --- | --- | --- | --- | --- | --- |
| Alcohols | 1 | - | 0.31 (0.578) | - | 0.68 (0.425) |
| Aldehydes | 1 | 0.15 (0.703) | 0.04 (0.850) | 0.05 (0.832) | 0.00 (0.953) |
| Alkanes | 1 | 0.23 (0.630) | 0.29 (0.588) | 0.97 (0.342) | 0.47 (0.503) |
| Benzene<br>Derivatives | 1 | 2.42 (0.119) | 0.05 (0.820) | 1.00 (0.334) | 0.25 (0.624) |
| Esters | 1 | - | 0.40 (0.529) | - | 0.68 (0.424) |
| Ketones | 1 | 1.02 (0.311) | 2.89 (0.089) | 0.02 (0.884) | 0.55 (0.473) |
| Monoterpenoids | 1 | - | 0.14 (0.713) | - | 0.03 (0.861) |
| Sesquiterpenoids | 1 | 0.26 (0.612) | 0.12 (0.725) | 0.05 (0.834) | 0.61 (0.448) |
| TotalVOC | 1 | 0.27 (0.606) | 0.35 (0.557) | 0.05 (0.827) | 0.39 (0.541) |
| MT_class | 3 | - | 5.14 (0.162) | - | 0.59 (0.633) |
| SQT_class | 3 | 7.64 (0.054) | **10.68 (0.014)\*** | 0.68 (0.580) | 1.05 (0.410) |
| Treatment | 3 | **11.09 (0.011)\*** | 5.14 (0.162) | 2.19 (0.142) | **4.84 (0.022)\*** |

